## Supplementary Information for "Evolutionary transition from a single RNA replicator to a multiple replicator network"

**This file includes:**

Supplementary Text

Figs. S1 to S17

Tables S1 to S3

References

### Supplementary Text

#### 1. Two additional long-term replication experiments

We performed two additional long-term replication experiments (E2 and E3). Initiated with the droplet mixture at round 76 of the main experiment (Fig. 1c), we performed independent 164 rounds (total 240 rounds) of serial transfer in each experiment. Host RNA concentrations showed similar yet distinct oscillation patterns (Supplementary Figs. S5a and b). The size of dominant parasitic RNAs for the additional experiments was similar to that for the main long-term replication experiment. We then performed PacBio sequencing for host and ~500 nt parasitic RNAs (if detected) at 12 points between rounds 92 and 239 of E2 and 9 points between rounds 114 and 239 of E3. From 4270–10000 reads of the host and parasitic RNAs (Supplementary Table S1), we identified 79 and 37, and 77 and 30 dominant mutations in host and parasitic RNAs for E2 and E3, respectively. Among these mutations, 30 and 20 mutations in host RNAs of E2 and E3, respectively, and 17 and 10 mutations in parasitic RNAs of E2 and E3, respectively, were not found in the dominant mutations of the main experiment, although 42 and 20 mutations were common in host and parasitic RNAs in all long-term replication experiments (Supplementary Figs. S6a and b). These data show that different host and parasitic RNAs evolved in the different experiments.

Next, we created consensus genotypes and phylogenetic trees based on the dominant mutations identified in both experiments (Supplementary Figs. S6c and d). We first defined the ancestral host RNA lineages (HL0) so that these lineages include all genotypes that have the same sets of mutations found in the ancestral host RNA lineage in the main experiment. In addition, we defined two host RNA lineages (HL4 and HL5) and two parasitic RNA lineages (PL4 and PL5) in E2, and three host RNA lineages (HL6, HL7, and HL8) and two parasitic RNA lineages (PL6 and PL7) in E3, which accumulated different sets of mutations. Although the phylogenetic trees indicate relatedness between PL4 and PL5 and between PL6 and PL7, these parasitic RNA lineages had different deletion regions and may have originated from distinct host RNAs, as described in Supplementary Text 3.

We then analyzed the population dynamics of each lineage using the 100 most frequent host and parasitic RNA genotypes at each sequenced round. The genotypes in different lineages show distinct patterns of mutation accumulation (Supplementary Figs. S8 and S9). The frequency of each lineage for both E2 and E3 (Supplementary Fig. S10) throughout the rounds revealed similar trends to the main long-term replication experiment (Figs. 3 and c), i.e., gradual diversification toward relatively stable coexistence of the evolved lineages. In E2, the host RNA lineages, HL5 and HL6, were first detected at round 33. Their frequencies varied from less than 0.1% to nearly 100% up to round 200 and afterwards were persistently maintained at more than 15% of the population. In parasitic RNAs, PL4 was detected throughout the sequenced rounds with more than 46% of the population, but PL5 coexisted as a dominant lineage from round 200 with similar frequencies. In E3, the host RNA lineages, HL8, HL9, and HL10 successively appeared. Their frequencies varied from less than 0.1% to nearly 100% up to round 198, and thereafter, all three lineages were consistently maintained at more than 0.5% of the population. In parasitic RNAs, PL6 was detected throughout the sequenced rounds with more than 83% of the population, but PL5 coexisted as 0.1–17% of the population from round 198. Overall, the four and five RNA lineages in E2 and E3, respectively, coexisted in the last ~40 rounds. These results support the possibility that the self-replicating RNA complexifies toward replicator communities through Darwinian evolution.

### 2. Investigation of higher-order interactions between the selected RNA clones

The interactions between RNA clones (Fig. 3d) were determined by examining each RNA replication by a specific RNA. However, in the long-term replication experiment (Fig. 1c), more than two types of RNAs were co-replicated, which may show more complex interactions. To examine the existence of such higher-order interactions, we incubated each RNA clone, a pair of two RNA clones, or a combination of three RNA clones for all possible RNA sets of selected rounds at 37 °C for 5 h and determined the extent of RNA (co-) replications through the translation of replicases (Supplementary Fig. S13). Next, we estimated the contribution of higher-order interactions arising only in the presence of three RNA clones by subjecting the replication data to Bahadur expansion analysis[1,2]. In Bahadur expansion analysis, the measured replication amounts of each RNA were converted into an orthogonal system consisting of interaction terms. For example, considering interactions among $RNA_i$, $RNA_j$, and $RNA_k$, the contributions of $RNA_j$, $RNA_k$, and a set of $RNA_j$ and $RNA_k$ to the replication of $RNA_i$ were quantified as Bahadur coefficients $w_j$, $w_k$, and $w_{jk}$, respectively, in a comparable form (lower coefficients indicate smaller contributions, and vice versa). Supplementary Fig. S14 shows the calculated Bahadur coefficients in all examined combinations of three RNA clones and the sum of coefficients of determination ($R^2$) for first-order Bahadur coefficients (e.g., $w_j$ and $w_k$), which represent the contribution of direct interactions between two RNAs. $R^2$ values in 40 out of 45 cases exceeded 0.7; in the other five cases, the effect of measurement errors became significant because all Bahadur coefficients were extremely low. These results indicated that higher-order interactions among three RNA clones did not significantly contribute to our results; the interdependent replication of RNA clones can essentially be understood from interactions between two RNAs, as represented in Fig. 3d.

### 3. Deletion sites of dominant parasitic RNAs

For parasitic RNA lineages that appeared in the main long-term replication experiment, PL1 deleted 224–536 and 744–1961 nt based on the ancestral host RNA sequence, as observed for parasite-γ in the previous study[3], whereas PL2 and PL3 deleted 178–526 and 766–1943 nt, which was not previously identified. For parasitic RNA lineages in the two additional experiments, PL5 and PL7 deleted similar RNA regions to PL2 and PL3, whereas PL4 and PL6 deleted previously unidentified regions, 173–293 and 673–1956 nt, and 224–536 and 745–1955 nt, respectively.

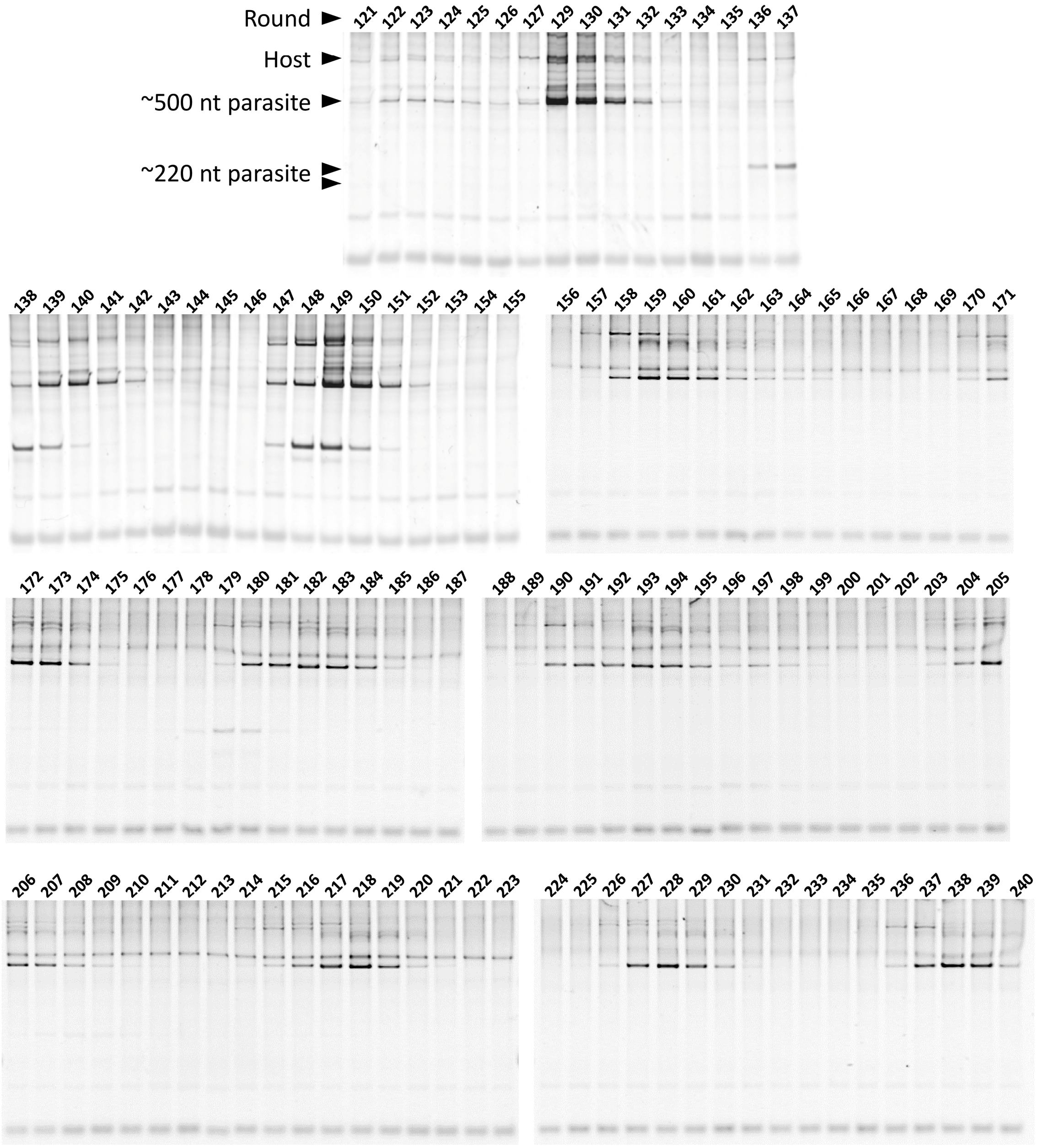


**Fig. S1 | Native polyacrylamide gel analysis of parasitic RNAs during the long-term replication experiment.** RNA mixtures at rounds 121–240 were subjected to native polyacrylamide gel electrophoresis. Band intensities of ~220 nt and ~500 nt parasitic RNAs were quantified and plotted in Fig. 1c. Multiple bands were sometimes detected in each class of the parasitic RNAs due to structural or size heterogeneity.

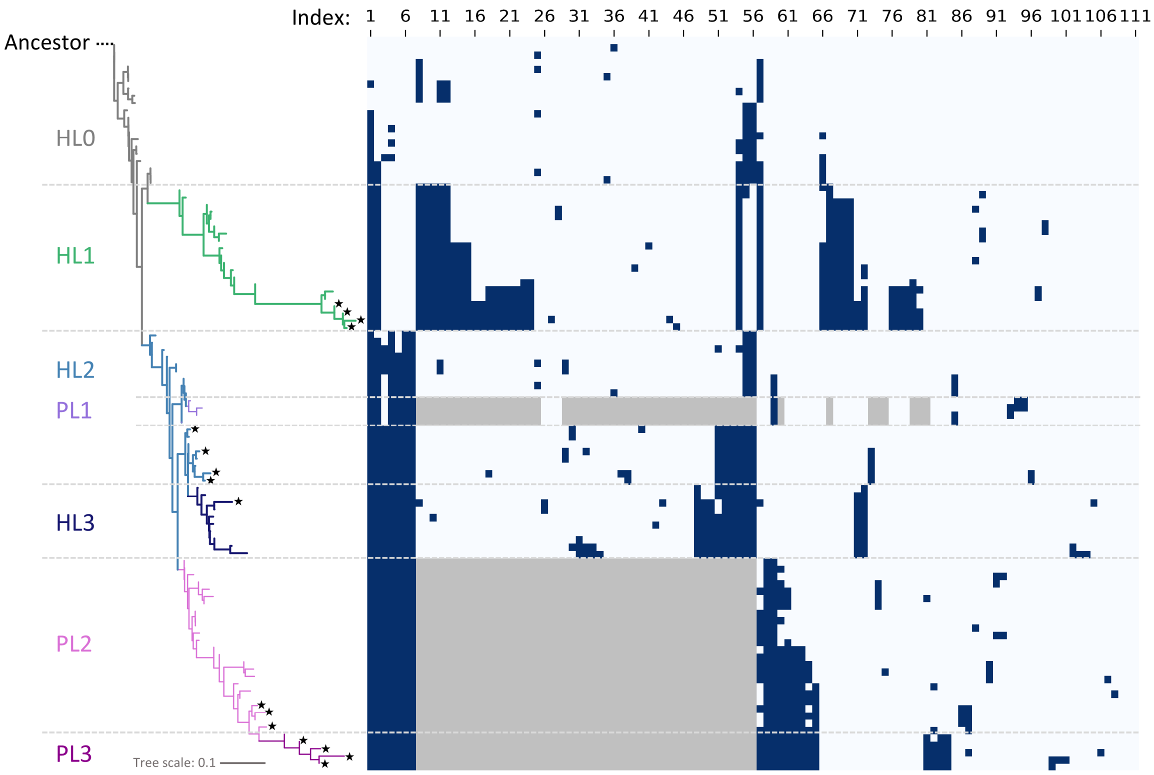


**Fig. S2 | An enlarged view of the dominant mutation map in Fig. 2.** Navy and grey colors indicate the presence of a point mutation and deletion, respectively. Mutation indices at the top correspond to ones in Supplementary Figs. S3 and S4.

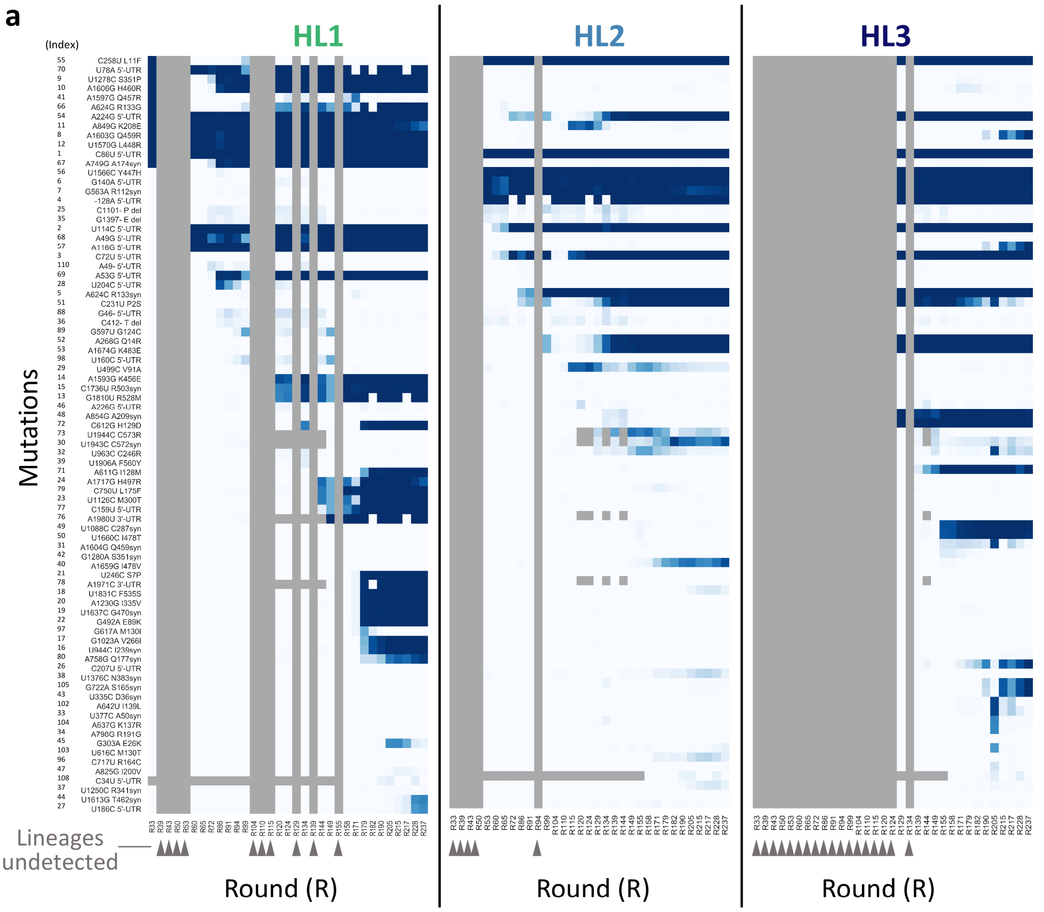


b

| | |
|---|---|
| only HL1 | 28 |
| only HL2 | 3 |
| only HL3 | 6 |
| only HL1 & 2 | 0 |
| only HL1 & 3 | 4 |
| only HL2 & 3 | 9 |
| All three lineages | 3 |

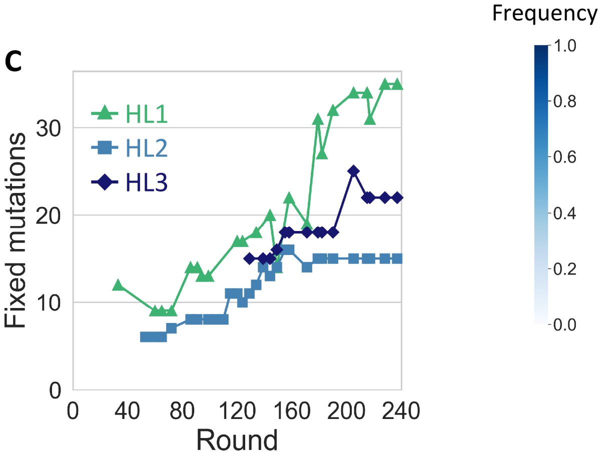


**Fig. S3 | Dominant mutations and fixation dynamics in the host RNA lineages. a,** Dominant mutations accumulated in HL1, HL2, and HL3 over rounds. The base numbers are based on the original host RNA. "syn" and "del" in mutation names stand for synonymous and deletion, respectively. Numbers to the left of the mutation names correspond to mutation indices shown in Supplementary Fig. 2. The intensity of the blue color indicates fixation frequency. Lineages were not detected at rounds marked with grey arrowheads. Some mutations near 5′ and 3′ ends (e.g., index 73, U1994C, C573R) were not determined at several rounds because we used different primers for efficient cDNA library preparation. **b**, Number of fixed mutations (accumulated in more than 50% sequences) in specific lineages at the last sequenced round (237). **c**, Number of fixed mutations in each lineage over rounds.

**a**

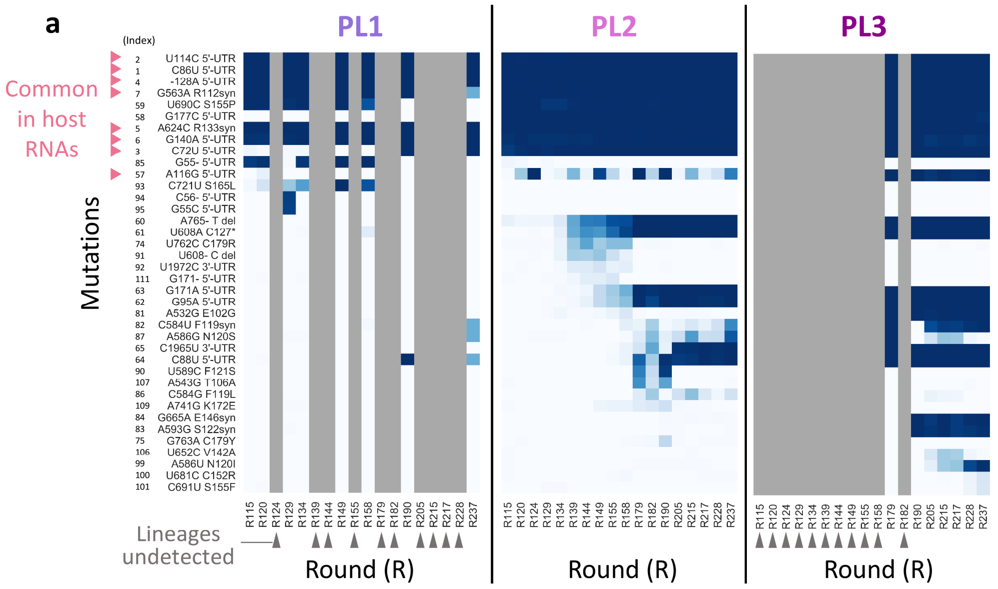


**b**

| | |
|---|---|
| only PL1 | 0 |
| only PL2 | 1 |
| only PL3 | 4 |
| only PL1 & 2 | 0 |
| only PL1 & 3 | 0 |
| only PL2 & 3 | 11 |
| All three lineages | 6 |

**c**

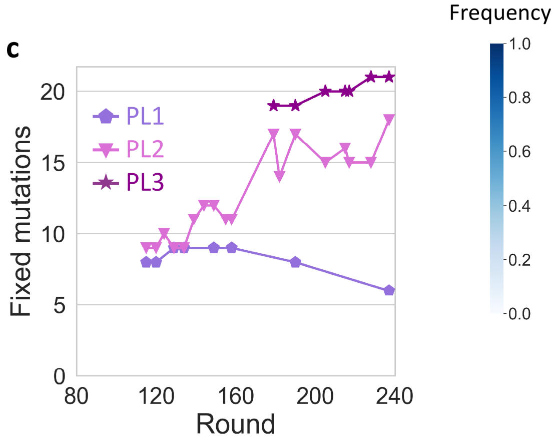


**Fig. S4 | Dominant mutations and fixation dynamics in the parasitic RNA lineages.** **a**, Dominant mutations accumulated in PL1, PL2, and PL3 over rounds. The base numbers are based on the original host RNA. "syn" and "del" in mutation names stand for synonymous and deletion, respectively. Numbers to the left of the mutation names correspond to mutation indices shown in Supplementary Fig. S2. The intensity of the blue color indicates fixation frequency. Grey regions indicate that mutations were not observed because lineages were not detected (indicated with grey arrowheads at rounds). Pink arrowheads indicate mutations commonly observed in host RNAs (Supplementary Fig. S3). Deletions at recombination sites were not shown. **b**, The number of fixed mutations (accumulated in more than 50% sequences) in specific lineages at the last sequenced round (237). Deletions due to different recombination sites were not counted. **c**, Number of fixed mutations in each lineage over rounds.

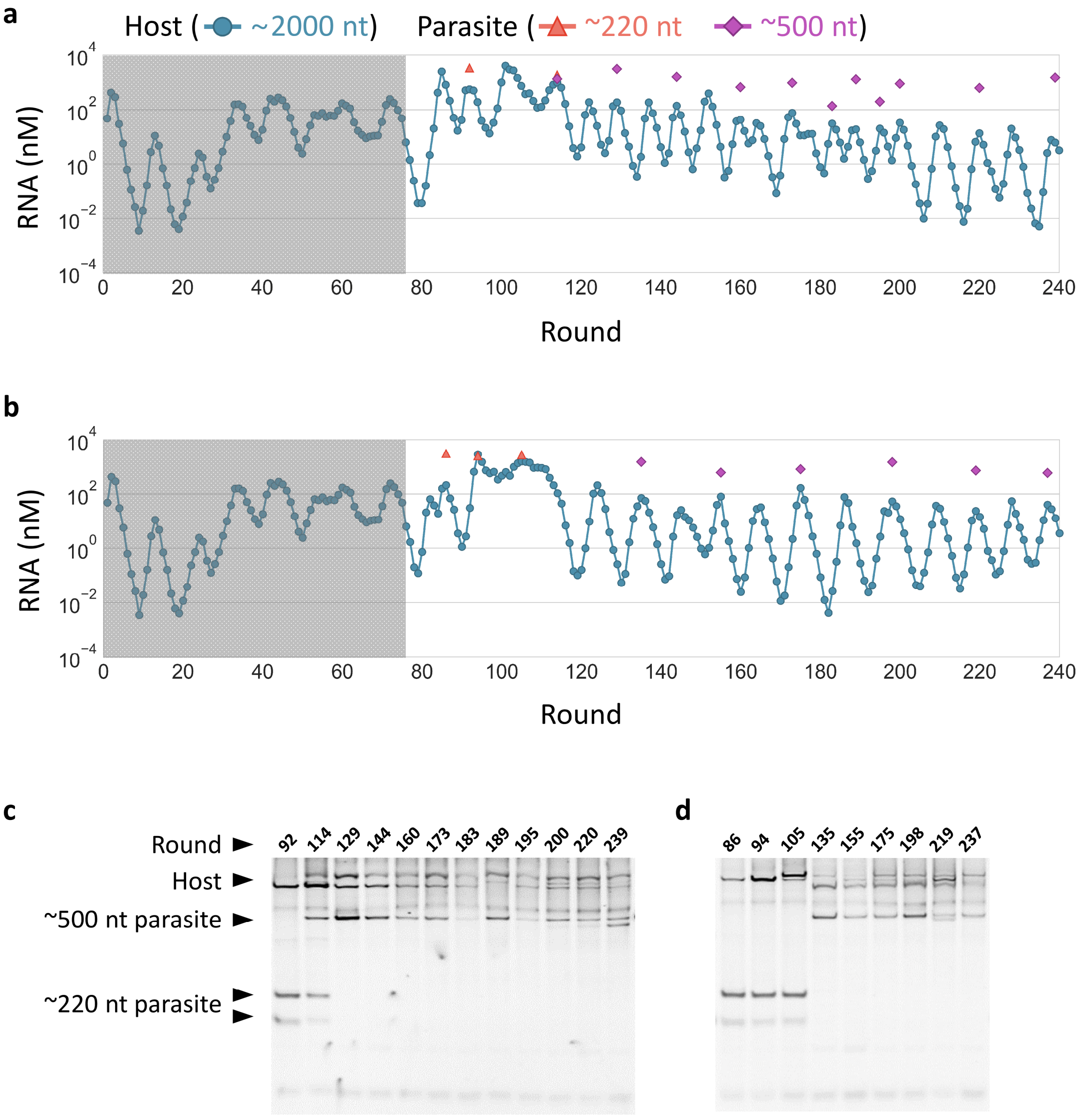


**Fig. S5 | Two additional long-term replication experiments. a–b**, Concentration changes of host and parasitic RNAs of different lengths in E2 (a) and E3 (b), where we newly performed 164 cycles of replications started with the droplet mixture at round 76 of the main long-term replication experiment. Parasitic RNA concentrations were determined only at sequenced rounds. The plot of host RNA concentrations in the shaded regions (up to 76 round) is the same as that of Fig. 1c. **c–d**, Native polyacrylamide gel electrophoresis of RNA mixtures in E2 (c) and E3 (d). Band intensities of ~220 nt and ~500 nt parasitic RNAs were quantified and plotted in panels a and b. Multiple bands were sometimes detected in each class of the parasitic RNAs due to structural or size heterogeneity.

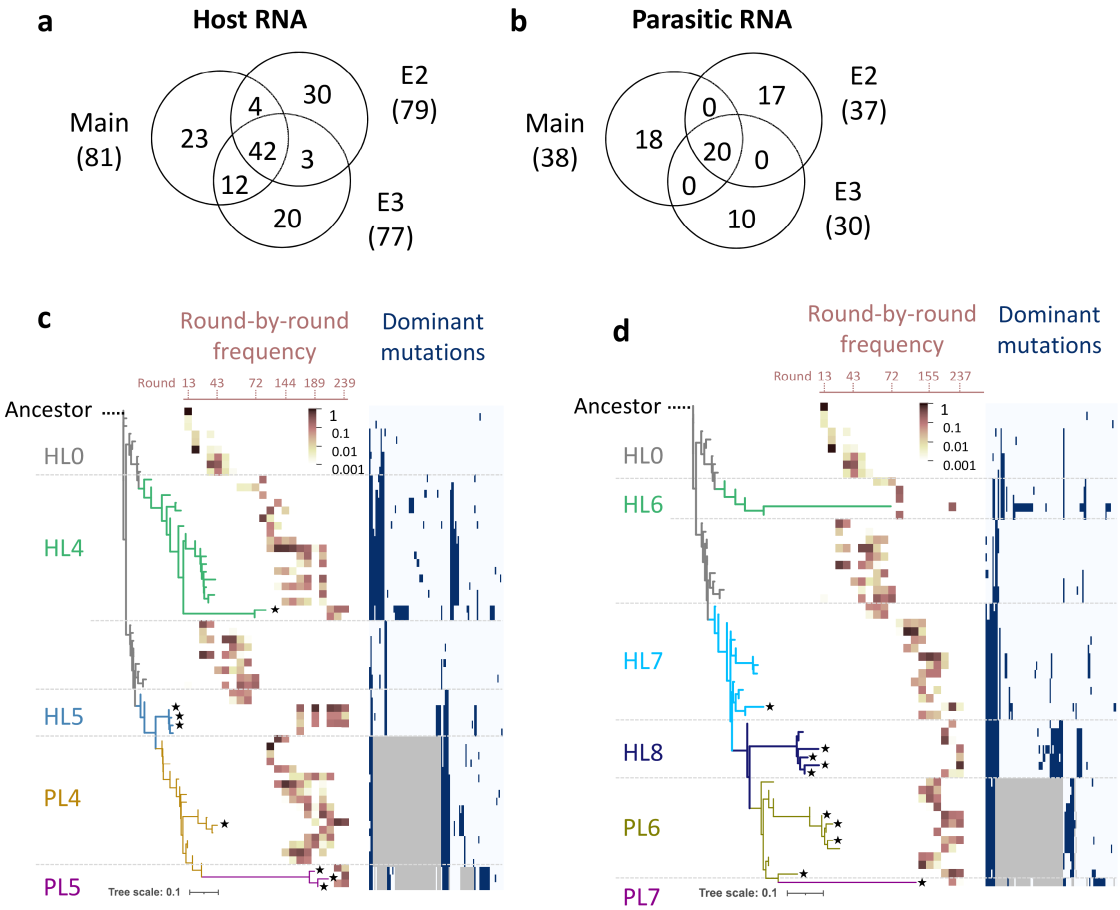


**Fig. S6 | Sequence and phylogenetic analyses of the two additional long-term replication experiments. a–b**, Venn diagrams showing the number of dominant mutations in host RNAs (a) and parasitic RNAs (b) identified in each of the three long-term replication experiments. Numbers in parenthesis at the name of experiments indicate the total mutation numbers. **c–d**, Phylogenetic trees were constructed based on the three most frequent host and parasitic RNA genotypes in all sequenced rounds for E2 (c) and E3 (d). The ancestral host RNA ("Ancestor") was designated as the root of the trees. Branches comprising defined lineages are colored differently. Host and parasitic RNA lineages are shown as thick and thin lines, respectively. The heatmaps superimposed on the trees show the frequencies of each genotype over all sequenced rounds (from left to right). Black star shapes at the tips of branches mark genotypes that remained to the last sequenced round. The lists of dominant mutations are shown on the right; navy and grey colors indicate the presence of a point mutation and deletion, respectively. An enlarged view of the list for each experiment is available in Supplementary Fig. S7.

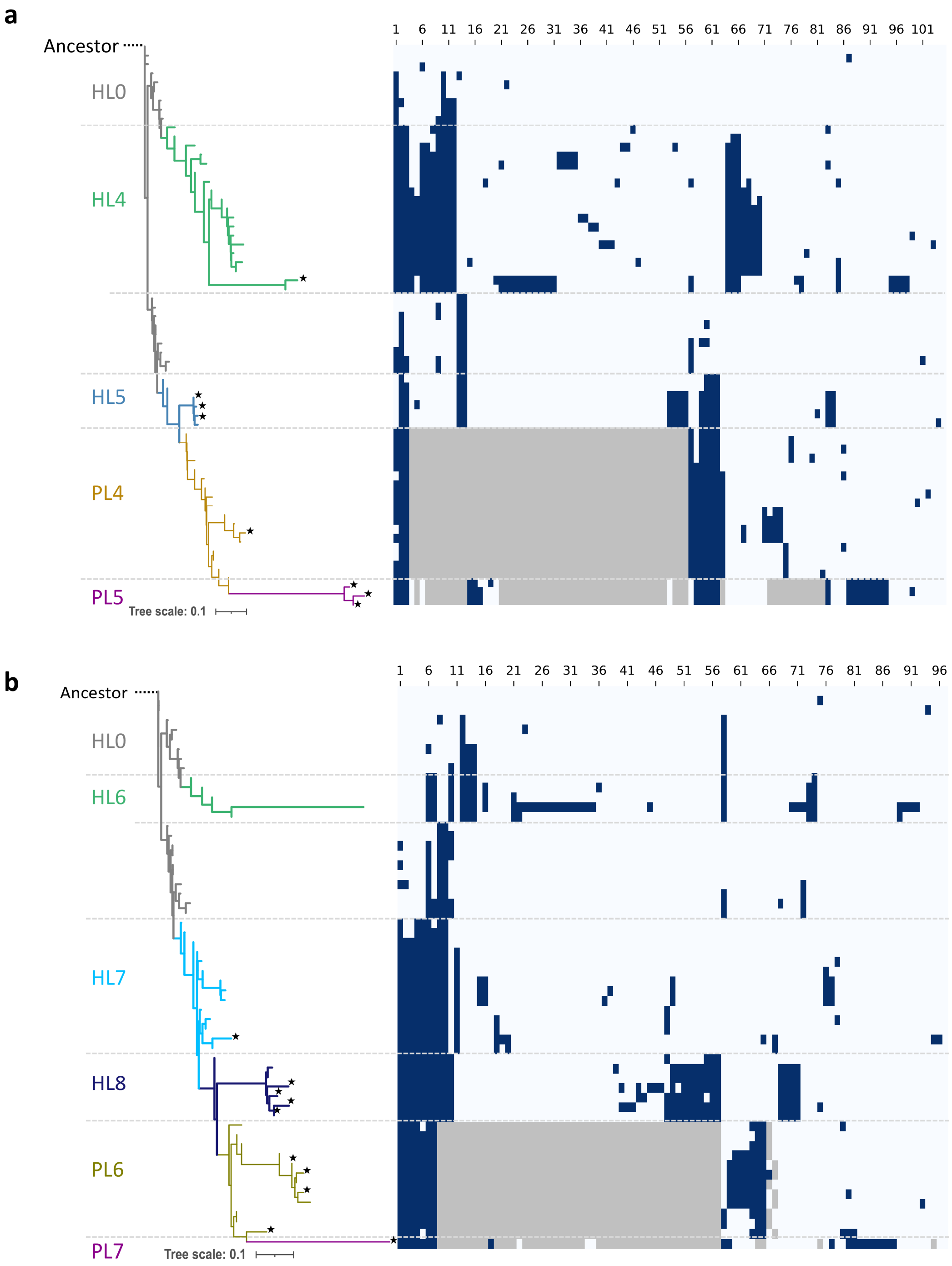


**Fig. S7. Enlarged views of the dominant mutation maps in Supplementary Figs. S6c and d. a–b**, The maps correspond to those in Supplementary Figs. S6c (a) and d (b). Navy and grey colors indicate the presence of a point mutation and deletion, respectively. Mutation indices at the top correspond to ones in Supplementary Figs. S8 (a) and S9 (b).

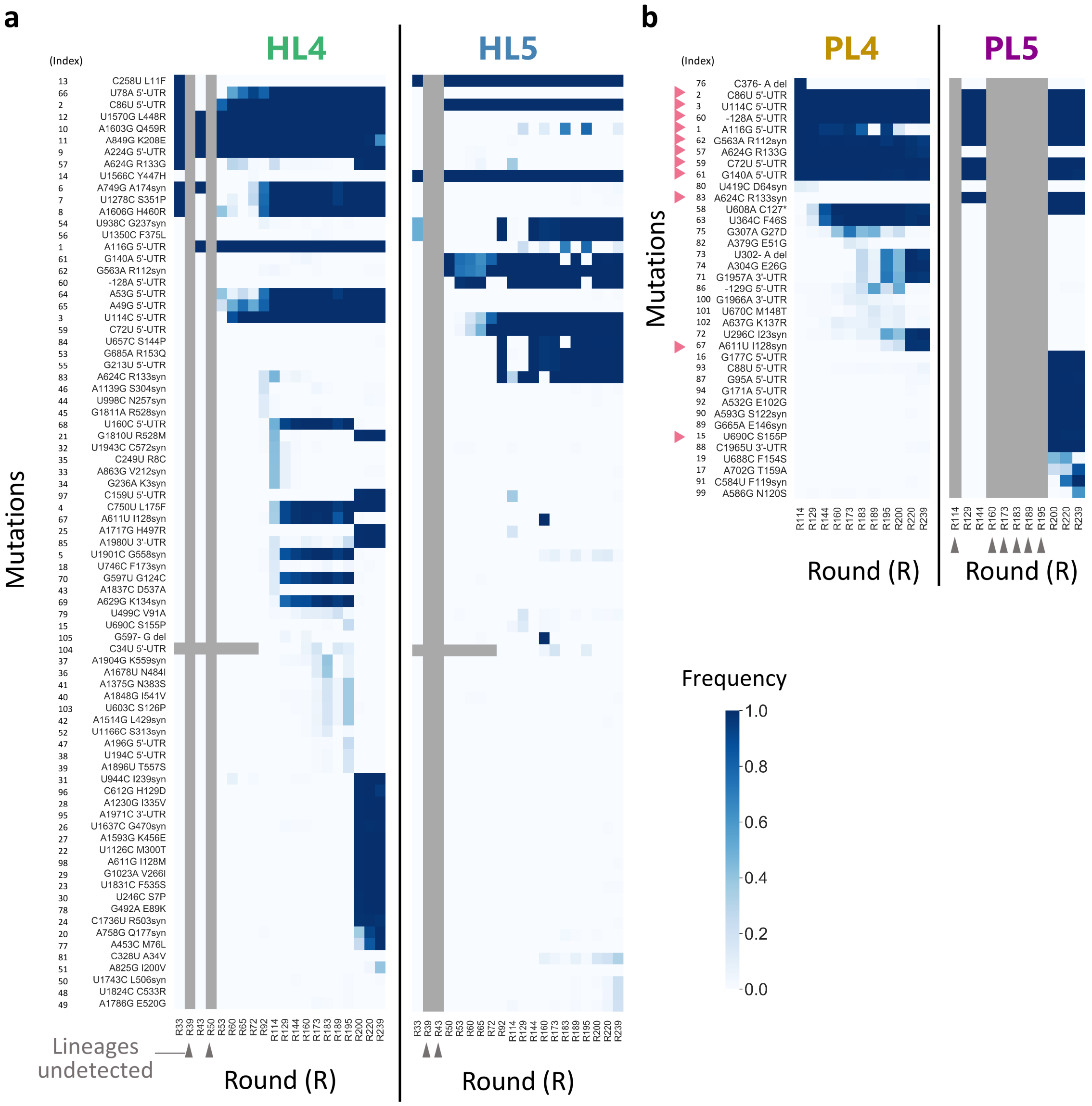


**Fig. S8 | Dominant mutations and fixation dynamics in the host and parasitic RNA lineages in E2. a–b**, Dominant mutations accumulated in HL4 and HL5 (a), or PL4 and PL5 (b) over rounds. The base numbers are based on the original host RNA. "syn" and "del" in mutation names stand for synonymous and deletion, respectively. Numbers to the left of the mutation names correspond to mutation indices shown in Supplementary Fig. S7. The intensity of the blue color indicates fixation frequency. Grey regions indicate that mutations were not observed because lineages were not detected (indicated with grey arrowheads at rounds) or different primers were used for cDNA library preparation. Pink arrowheads at mutations for parasitic RNA lineages indicate those commonly observed in host RNAs

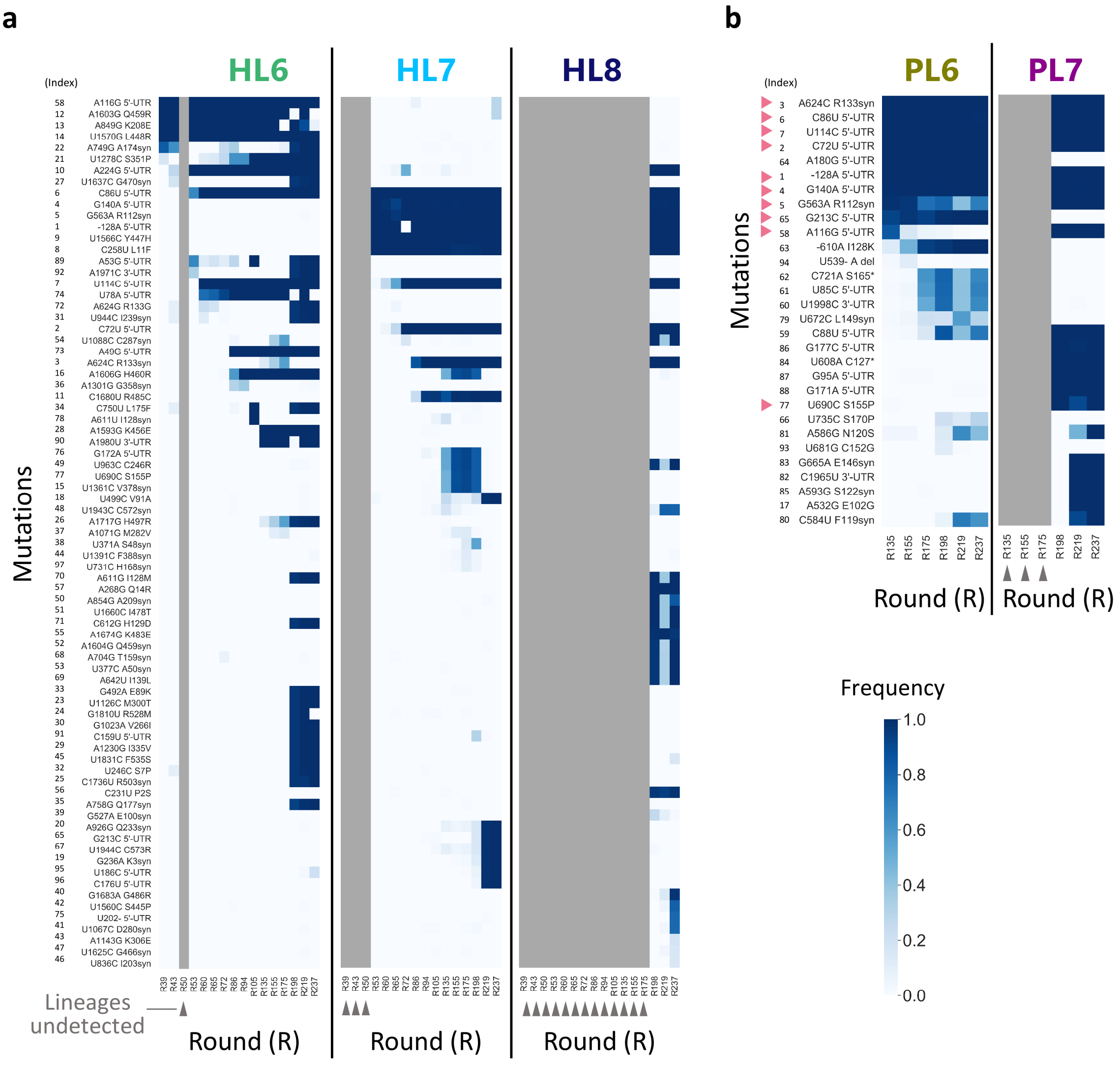


**Fig. S9. Dominant mutations and fixation dynamics in the host and parasitic RNA lineages in E3. a–b**, Dominant mutations accumulated in HL6, HL7, and HL8 (a), or PL6 and PL7 (b) over rounds. The base numbers are based on the original host RNA. "syn" and "del" in mutation names stand for synonymous and deletion, respectively. Numbers to the left of the mutation names correspond to mutation indices shown in Supplementary Fig. S7. The intensity of the blue color indicates fixation frequency. Grey regions indicate that mutations were not observed because lineages were not detected (indicated with grey arrowheads at rounds). Pink arrowheads at mutations for parasitic RNA lineages indicate those commonly observed in host RNAs.

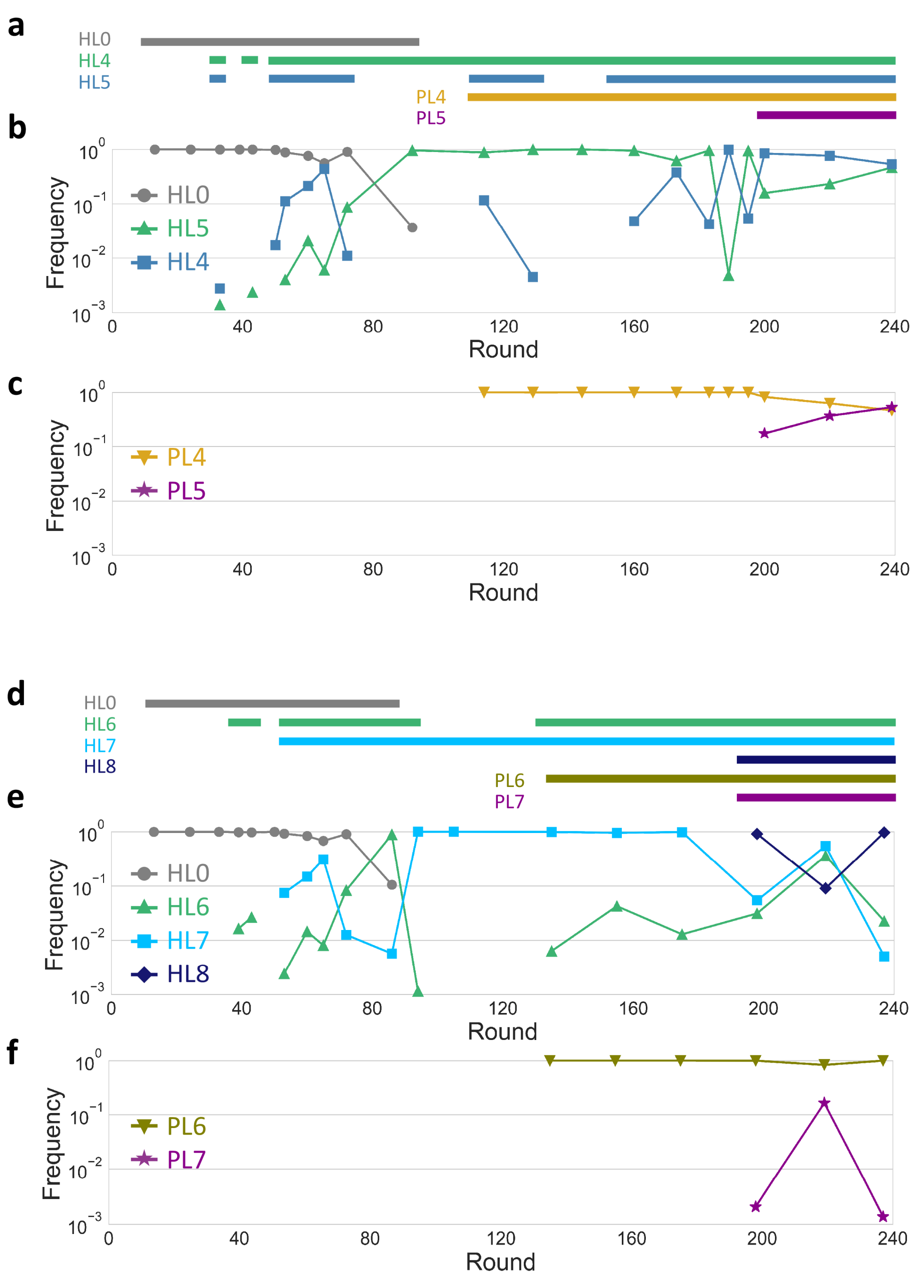


**Fig. S10 | Population dynamics of the lineages in the additional long-term replication experiments. a–f**, Frequencies of the lineages in total sequence reads of the analyzed genotypes for host (b for E2 and e for E3) and parasitic (c for E2 and f for E3) RNAs. Horizontal lines above the graphs (a, d) indicate rounds where the frequency of each lineage in the same color was plotted (above 0.1 %).

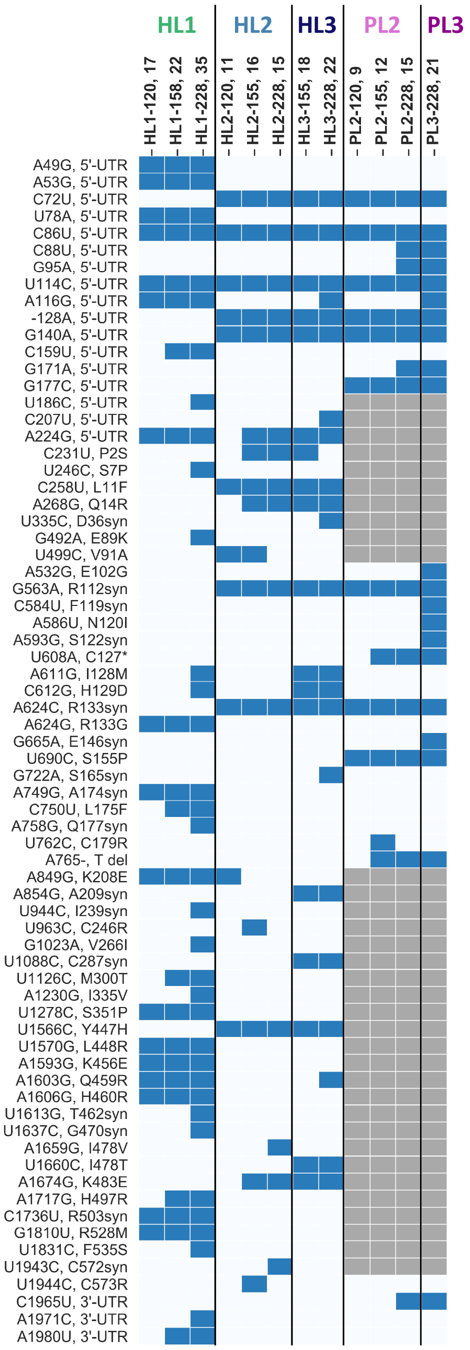

**Fig. S11 | List of mutations in the selected RNA clones.** The base numbers are based on the original host RNA. “syn” and “del” in mutation names stand for synonymous and deletion, respectively. Grey regions indicate deleted sites. The numbers after the names of each clone show mutation numbers.

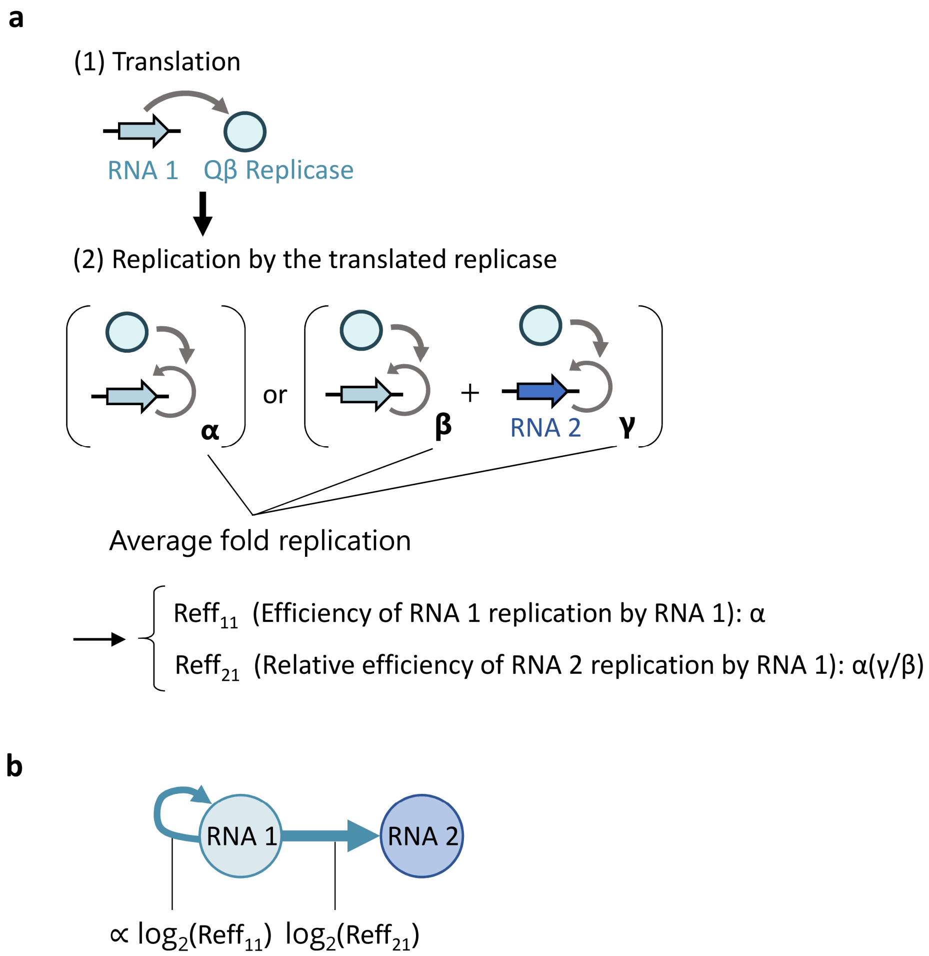


**Fig. S12 | Procedure for the depiction of replication relationships between the RNA clones by directed graphs. a**, In a translation-uncoupled replication experiment, where the translation of the replicase gene from RNA 1 was followed by RNA replication in the presence or absence of RNA 2 (Fig. 4a), three types of fold replications, RNA 1 "self" replication in the absence of RNA 2 (α), RNA 1 replication in the presence of RNAs 1 and 2 (β), and RNA 2 replication in the presence of RNAs 1 and 2 (γ), were determined. Using the average of these replications above a background level (>1.5-fold), the efficiencies of RNA 1 and RNA 2 replications by the replicase translated from RNA1, $Reff_{11}$ and $Reff_{21}$, were determined. **b**, Directed graphs were depicted by setting the widths of arrows proportional to the binary logarithm of $Reff_{11}$ and $Reff_{21}$.

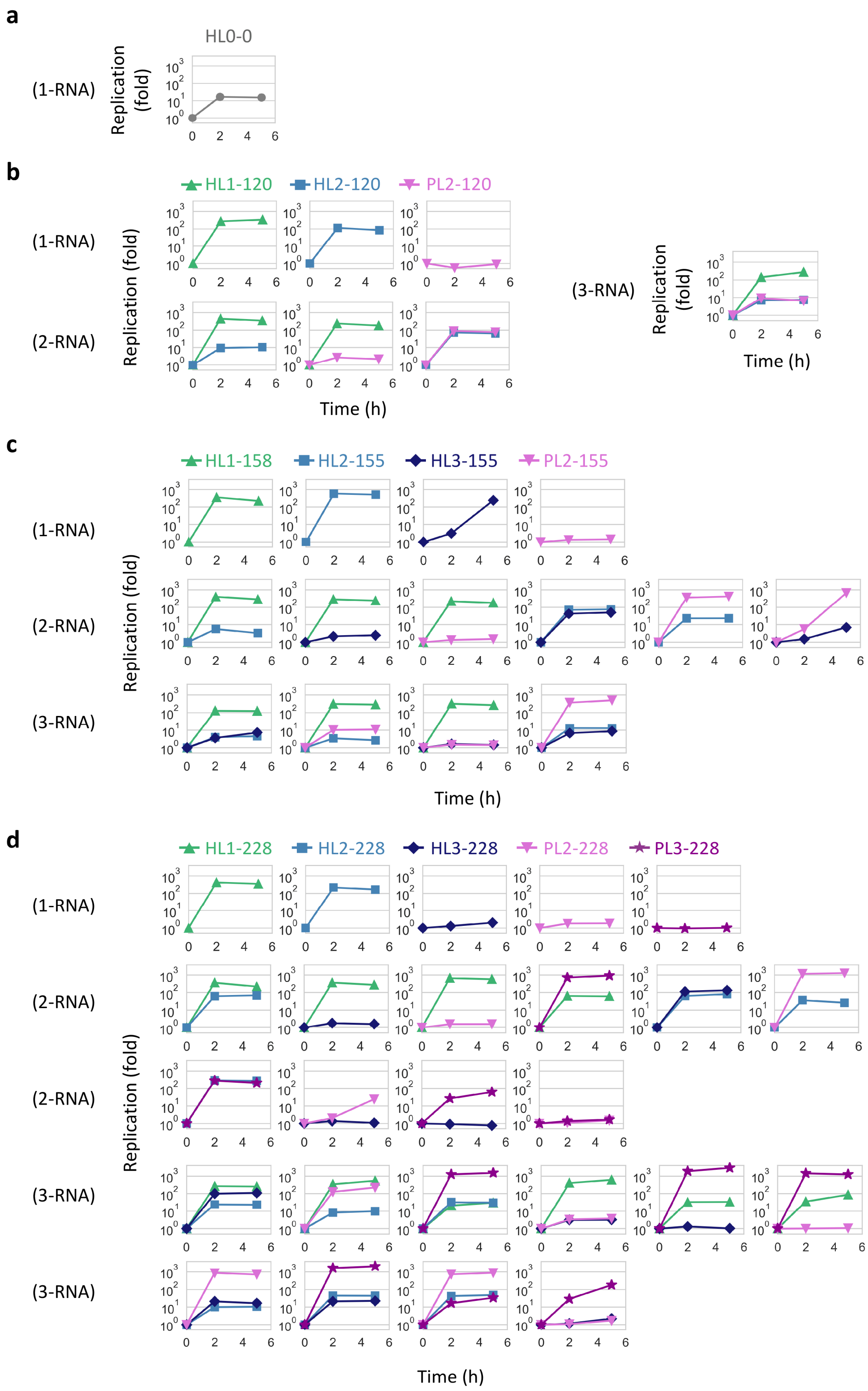


**Fig. S13 | Translation-coupled replication experiments. a–d**, One, two, or three RNA clones (10 nM each) at rounds 0 (a), 120 (b), 155–158 (c), and 228 (d) were incubated at 37 °C for 5 h in the translation system, and replications of each RNA at 2 and 5 h were measured by sequence-specific RT-qPCR.

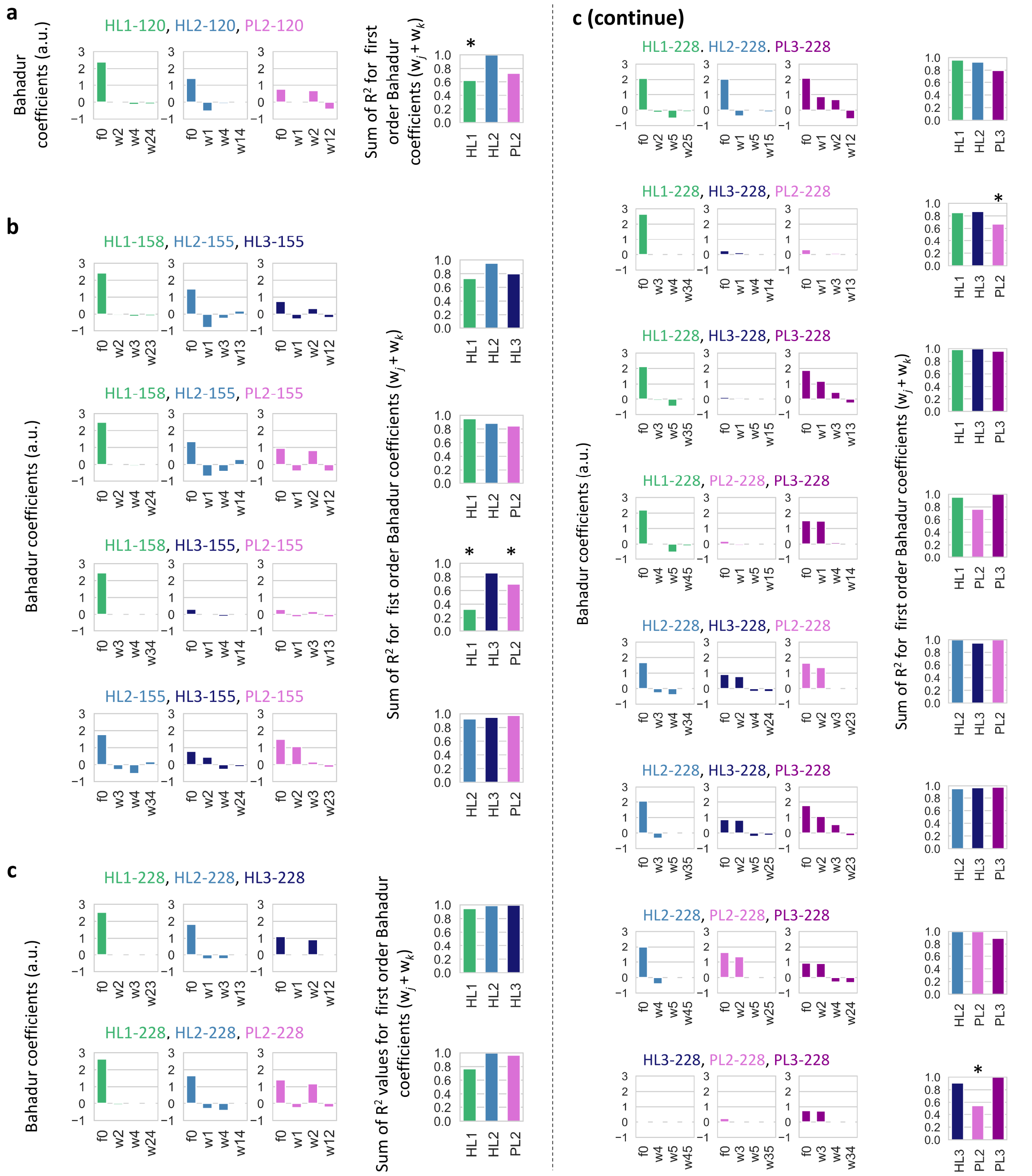


**Fig. S14 | Bahadur expansion analysis. a–c**, Bahadur coefficients (left) and the sum of coefficients of determination ($R^2$) for first order Bahadur coefficients (i.e., $w_j + w_k$) (right) for each combination of three RNA clones at rounds 120 (a), 155–158 (b), and 228 (c), calculated from fold replications (at 2 h) in the translation-coupled replication experiments (Supplementary Fig. S13). Number *j* in $w_j$ is 1, 2, 3, 4, and 5 for RNA clones in HL1, HL2, HL3, PL2, and PL3, respectively. 5 out of 45 cases for which calculated $R^2$ are low (<0.7) are indicated with asterisks.

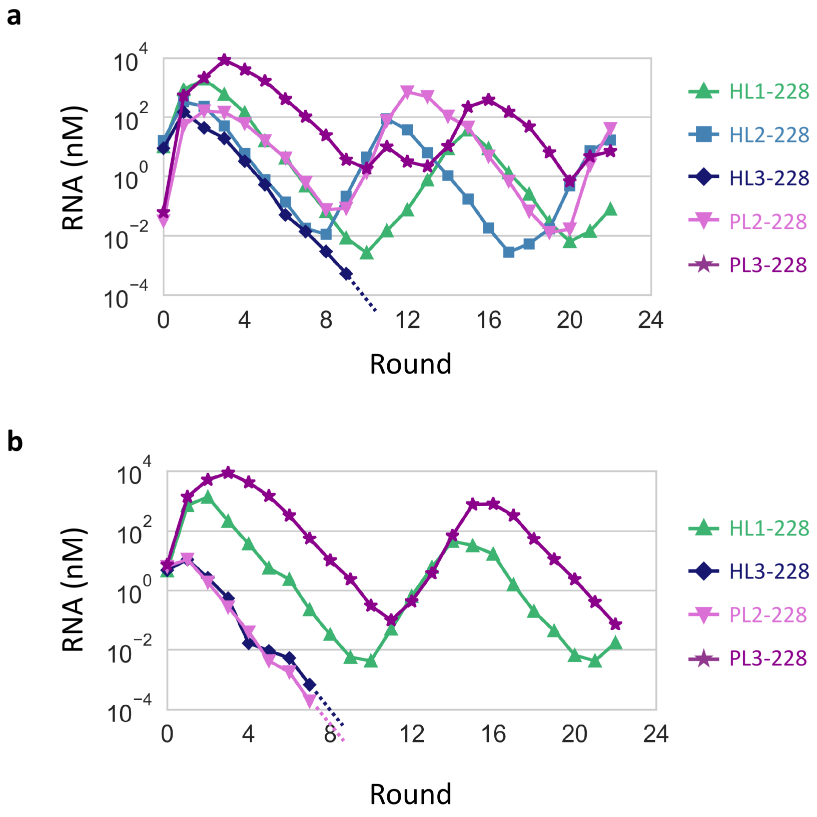


**Fig. S15 | Additional long-term replication experiments started with a mixture of the RNA clones at round 228. a–b**, RNA concentration changes in long-term replication experiments initiated with 10 nM each of HL1-, HL2-, and HL3-228, and 0.1 nM each of PL2- and PL3-228 (a) or 10 nM each of HL1-, HL3-, PL2-, and PL3-228 (b). The concentrations were measured by sequence-specific RT-qPCR.

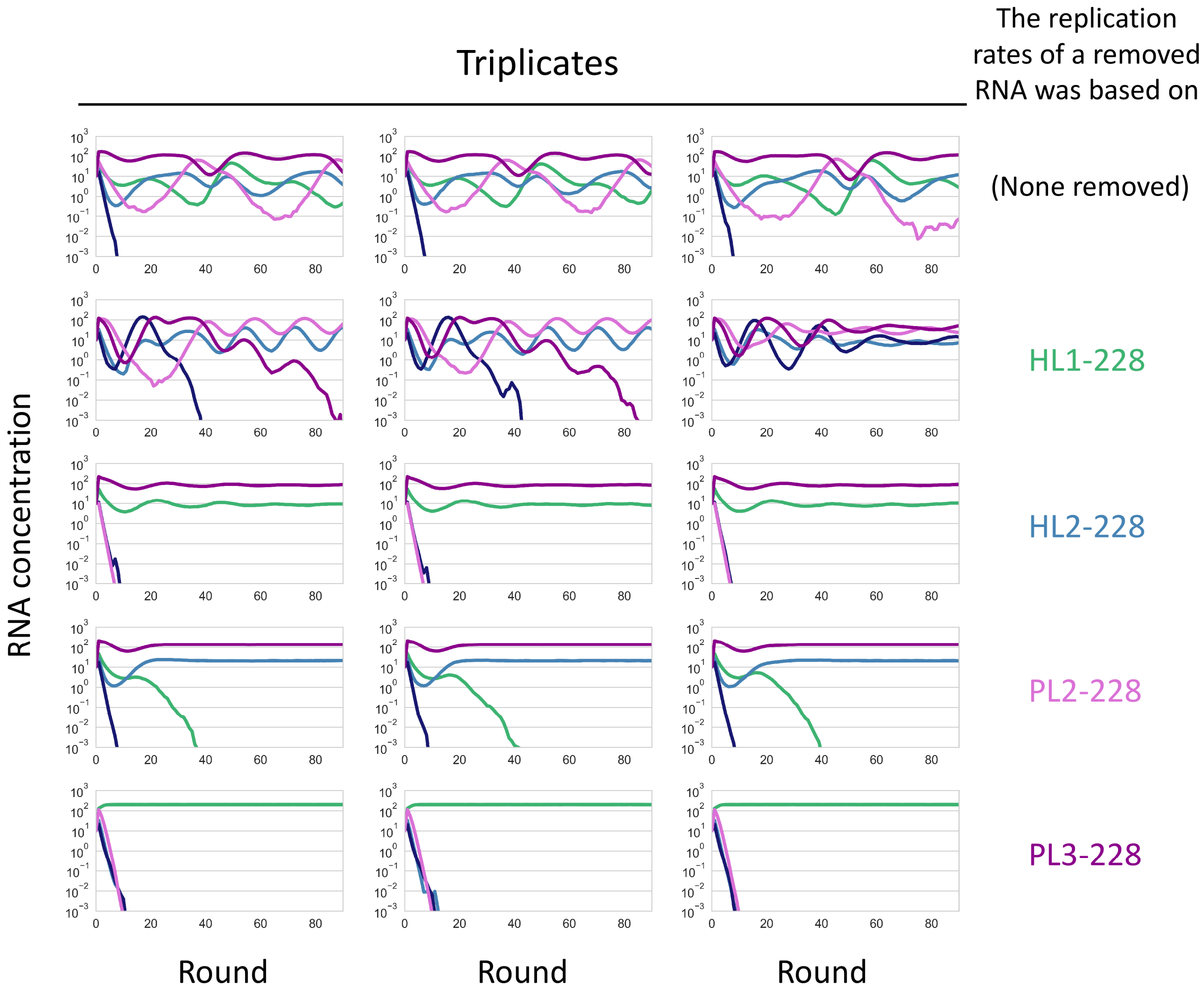


**Fig. S16 | Dynamics of the RNA replicator network in the absence of one of the RNAs.** Simulations were performed as that presented in Fig. 5b, in the absence of one of the four RNAs that sustainably replicated. Each simulation was performed three times independently. The upper leftmost panel is the same as Fig. 5b.

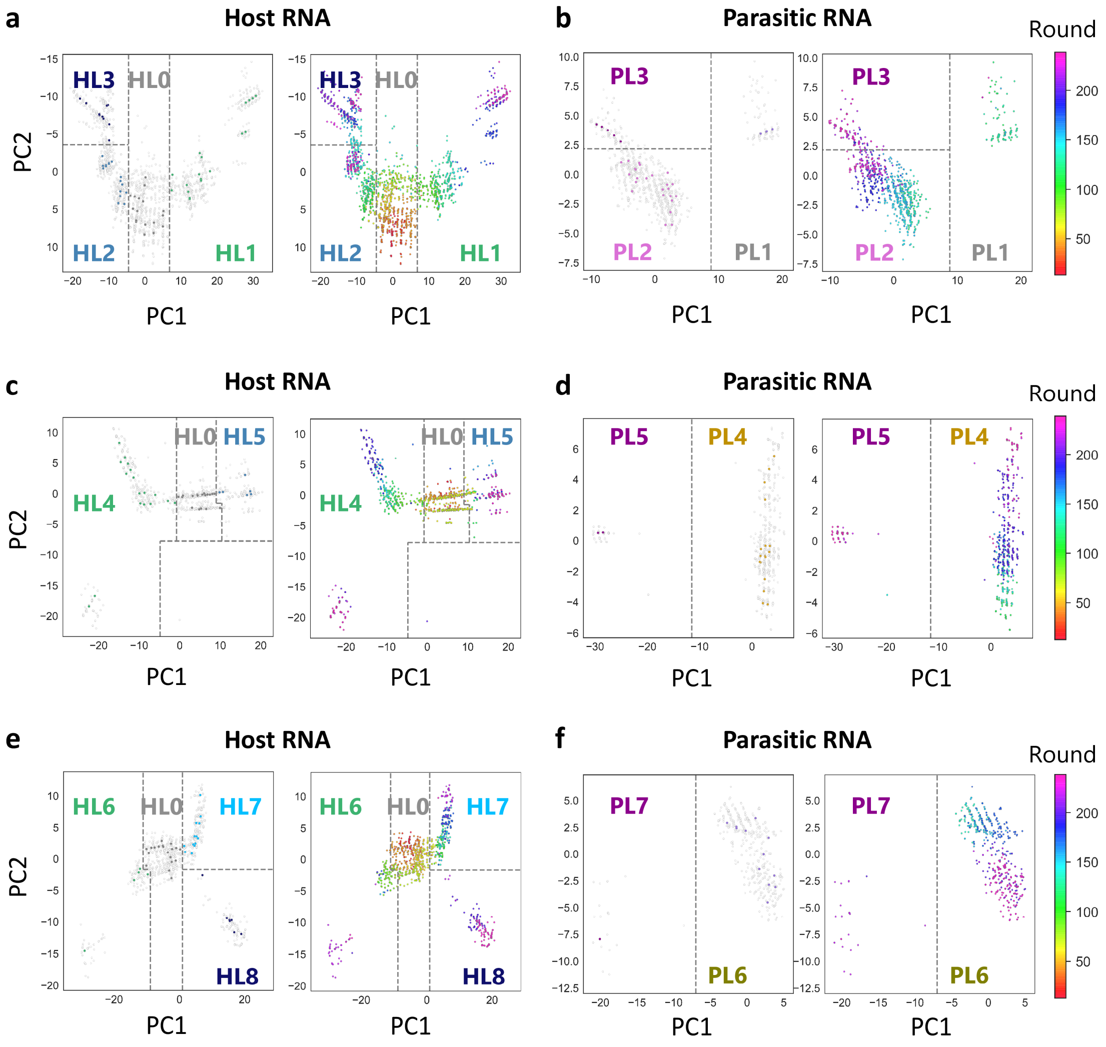


**Fig. S17 | Mapping of consensus host and parasitic RNA genotypes in sequence spaces. a–f**, Two-dimensional (2D) maps were created based on Hamming distances between all top 100 consensus host RNA genotypes or ~500 nt parasitic RNA genotypes obtained throughout the main long-term replication experiment (a, b), the additional long-term replication experiment E2 (c, d), and that of E3 (e, f). The Hamming distance matrices were plotted on the maps using Principal Coordinate Analysis for dimension reduction. In the 2D maps, each genotype is shown as a point; genotypes located in closer areas are expectedly more related. Left plots in each panel highlight genotypes represented in the phylogenetic trees (Fig. 2 and Supplementary Fig. S6), colored to indicate lineages defined based on the trees. All displayed genotypes were then classified in each lineage as indicated. Right plots show the same maps with genotypes colored to indicate the appearance of rounds.

**Table S1 | Number of analyzed reads obtained by PacBio sequencing.**

| Experiment | Spieces | Round | Read number |
|---|---|---|---|
| Main | Host | 13 | 4143 |
| Main | Host | 24 | 605 |
| Main | Host | 33 | 718 |
| Main | Host | 39 | 365 |
| Main | Host | 43 | 484 |
| Main | Host | 50 | 1020 |
| Main | Host | 53 | 1358 |
| Main | Host | 60 | 1409 |
| Main | Host | 65 | 3097 |
| Main | Host | 72 | 1364 |
| Main | Host | 86 | 2135 |
| Main | Host | 91 | 637 |
| Main | Host | 94 | 2058 |
| Main | Host | 99 | 855 |
| Main | Host | 104 | 2535 |
| Main | Host | 110 | 1758 |
| Main | Host | 115 | 4003 |
| Main | Host | 120 | 1202 |
| Main | Host | 124 | 1253 |
| Main | Host | 129 | 1879 |
| Main | Host | 134 | 841 |
| Main | Host | 139 | 2395 |
| Main | Host | 144 | 2329 |
| Main | Host | 149 | 1091 |
| Main | Host | 155 | 4998 |
| Main | Host | 158 | 10000 |
| Main | Host | 171 | 10000 |
| Main | Host | 179 | 10000 |
| Main | Host | 182 | 10000 |
| Main | Host | 190 | 10000 |
| Main | Host | 205 | 10000 |
| Main | Host | 215 | 10000 |
| Main | Host | 217 | 10000 |
| Main | Host | 228 | 9999 |
| Main | Host | 237 | 10000 |
| E2 | Host | 92 | 5882 |
| E2 | Host | 114 | 7840 |
| E2 | Host | 129 | 7259 |
| E2 | Host | 144 | 9056 |
| E2 | Host | 160 | 8477 |
| E2 | Host | 173 | 8415 |
| E2 | Host | 183 | 10000 |
| E2 | Host | 189 | 4270 |
| E2 | Host | 195 | 10000 |
| E2 | Host | 200 | 9288 |
| E2 | Host | 220 | 7209 |
| E2 | Host | 239 | 10000 |
| E3 | Host | 86 | 5393 |
| E3 | Host | 94 | 7149 |
| E3 | Host | 105 | 10000 |
| E3 | Host | 135 | 10000 |
| E3 | Host | 155 | 10000 |
| E3 | Host | 175 | 10000 |
| E3 | Host | 198 | 8910 |
| E3 | Host | 219 | 5187 |
| E3 | Host | 237 | 10000 |
| Main | Parasite | 115 | 1753 |
| Main | Parasite | 120 | 680 |
| Main | Parasite | 124 | 2073 |
| Main | Parasite | 129 | 1979 |
| Main | Parasite | 134 | 1102 |
| Main | Parasite | 139 | 1254 |
| Main | Parasite | 144 | 1099 |
| Main | Parasite | 149 | 758 |
| Main | Parasite | 155 | 840 |
| Main | Parasite | 158 | 10000 |
| Main | Parasite | 179 | 10000 |
| Main | Parasite | 182 | 10000 |
| Main | Parasite | 190 | 10000 |
| Main | Parasite | 205 | 9999 |
| Main | Parasite | 215 | 10000 |
| Main | Parasite | 217 | 10000 |
| Main | Parasite | 228 | 10000 |
| Main | Parasite | 237 | 10000 |
| E2 | Parasite | 114 | 5435 |
| E2 | Parasite | 129 | 8077 |
| E2 | Parasite | 144 | 9533 |
| E2 | Parasite | 160 | 7292 |
| E2 | Parasite | 173 | 8999 |
| E2 | Parasite | 183 | 7403 |
| E2 | Parasite | 189 | 4711 |
| E2 | Parasite | 195 | 4483 |
| E2 | Parasite | 200 | 5548 |
| E2 | Parasite | 220 | 7232 |
| E2 | Parasite | 239 | 8118 |
| E3 | Parasite | 135 | 10000 |
| E3 | Parasite | 155 | 10000 |
| E3 | Parasite | 175 | 10000 |
| E3 | Parasite | 198 | 9856 |
| E3 | Parasite | 219 | 4587 |
| E3 | Parasite | 237 | 10000 |

Grey regions indicate reads obtained in the previous study[3].

**Table S2 | Methods for the construction of each plasmid.**

| Evolved RNA clones encoded in plasmids | Methods of plasmid construction |
|---|---|
| HL1-120 | Site-specific mutagenesis of the plasmid encoding Host-99 in the previous study[3] |
| HL2-120 | Obtained in the previous study[3] |
| PL2-120 | Gene synthesis service of Eurofins Genomics |
| HL1-158 | Site-specific mutagenesis of the plasmid encoding Host-99 in the previous study[3] |
| HL2-155 | Site-specific mutagenesis of the plasmid encoding HL3-155 |
| HL3-155 | Gene synthesis service of Eurofins Genomics |
| PL2-155 | Site-specific mutagenesis of the plasmid encoding PL2-120 |
| HL1-228 | Site-specific mutagenesis of the plasmid encoding a randomly cloned RNA at round 190* |
| HL2-228 | Site-specific mutagenesis of the plasmid encoding HL3-155 |
| HL3-228 | Site-specific mutagenesis of the plasmid encoding HL3-155 |
| PL2-228 | Site-specific mutagenesis of the plasmid encoding PL2-155 |
| PL3-228 | Site-specific mutagenesis of the plasmid encoding PL2-155 |

*Cloning was performed by using SMARTer® RACE 5'/3' Kit (Takara) according to the manufacture's protocol.

*Cloning was performed by using SMARTer® RACE 5'/3' Kit (Takara) according to the manufacture's protocol.

### Table S3 | The list of primers (from 5' end to 3' end).

**Primers for the measurement of host RNA concentrations in the long-term replication experiments by quantitative RT-PCR**

| Names | Sequences | Description |
|---|---|---|
| Primer 1 | CAAGTATCGTAAGTTGCTGCC | Used in the main long-term replication experiment |
| Primer 2 | CCGTAATCACCGGTACGTAC | Used in the main long-term replication experiment |
| Primer 3 | GCTGCCTAAACAGCTGCAAC | Used in the additional long-term replication experiments (E2, E3) |
| Primer 4 | CGCTCTTGGTCCCTTGTATG | Used in the additional long-term replication experiments (E2, E3) |

**Primers for RT-PCR to prepare cDNA libraries for sequence analysis**

| Names | Sequences | Description (if any) |
|---|---|---|
| Primer 5 | CCCGGAAGGGGGGGACGAGG | |
| Primer 6 | GGGGGGTCACCTCGCGCAG | |
| Primer 7 | ACAACCCGAACAACAGCAC | Used instead of Primer 5 for host RNAs at round 120, 124, 130, 134 in the main long-term replication experiment |
| Primer 8 | GGGTCACCTCGCGCAGC | Used instead of Primer 6 for host and parasitic RNAs at round 120–155 samples in the main long-term replication experiment |

**Sequence specific primers for the detection of each RNA clone by quantitative RT-PCR**

| Names | Sequences | Target RNA clones |
|---|---|---|
| Primer 9 | ATACACATGGCTCGTAGAAAA | HL0-0 (Target RNA) |
| Primer 10 | GGCGTACACGCTTGCGGAAGT | HL0-0 |
| Primer 11 | CGAACGCTCGTCTCTATAGG | HL1-120 |
| Primer 12 | GTACACGCTTGCGGAAGC | HL1-120 |
| Primer 13 | AAGGTCGCGCCTCTCCA | HL2-120 |
| Primer 14 | ATGCTGTCTTAGGCATGTGT | HL2-120 |
| Primer 15 | AGGTCTCCGGCTGAATGTG | PL2-120 and PL2-155 |
| Primer 16 | CAAGCCTAACATACACGCTTG | PL2-120 |
| Primer 17 | TCCGTCCTTCAAGTTTGCGT | HL1-158 and HL1-228 |
| Primer 18 | CTTAGGTACGGTAACTGCTTC | HL1-158 and HL1-228 |
| Primer 19 | AAGGTCGCGCCTCTCCA | HL2-155 (with HL1-155) and HL2-228 |
| Primer 20 | CGCGAAGATGCTGTCTTAGA | HL2-155 (with HL1-155) and HL2-228 |
| Primer 21 | TGGGCGAGTCATGTATAC | HL2-155 (in the presence of both HL1-155 and HL3-155) |
| Primer 22 | CACGCTTGCGGAAGT | HL2-155 (in the presence of both HL1-155 and HL3-155) |
| Primer 23 | TACGCGATCGGTTGCGTC | HL2-155 (unless otherwise noted) |
| Primer 24 | TCGGGGCAATAAGAGCTCA | HL2-155 (unless otherwise noted) |
| Primer 25 | TGATATTAGCCCTTTTAATAAAGCG | HL3-155 |
| Primer 26 | TCGGGGCAATAAGAGCTCG | HL3-155 |
| Primer 27 | CAAGCCTAACAACGCGCTTG | PL2-155 |
| Primer 28 | AACAACGAATAACCGTTCA | HL3-228 |
| Primer 29 | TGTTCTTAGGTACGGTAACC | HL3-228 |
| Primer 30 | TCTAGAAAGTCTCCGGCTGA | PL2-228 |
| Primer 31 | GCAGTGACGCAACATATCC | PL2-228 |
| Primer 32 | TCTAGAAAGTCTCCGGCTGG | PL3-228 |
| Primer 33 | GCAGTGACGCAACATATCT | PL3-228 |
